# *Chlorobaculum tepidum* Outer Membrane Vesicles May Transport Biogenic Elemental Sulfur

**DOI:** 10.1101/2024.09.30.615447

**Authors:** Vadesse L. Noundou, Amalie Levy, Shannon Modla, Yanbao Yu, Jing Qu, Thomas E. Hanson

## Abstract

Outer membrane-derived vesicles (OMVs) have been studied in different phyla of Gramnegative bacteria, most extensively in the Pseudomonadota, where they have been shown to participate in diverse biological and environmental processes. To date, the production of OMVs has not been reported in the Chlorobiaceae within the phylum Chlorobiota. *Chlorobaculum. tepidum* is the model organism for the Chlorobiaceae that synthesizes and consumes insoluble extracellular sulfur (S(0)) globules by an unknown mechanism. Here, we report evidence implicating outer membrane vesicles in biogenic S(0) globule synthesis. We demonstrate that *Cba. tepidum* secretes OMVs in the extracellular milieu, and that OMV concentration and size vary with growth conditions, particularly sulfide concentration. A core of 31 proteins involved in diverse biological processes such as cell wall biogenesis, inorganic ion transport and metabolism were found to be shared between OMVs, extracellular S(0) globules and *Cba. tepidum* intact cells. Multiple analytical methods indicated that OMVs contain S(0) and that OMVs and biogenic S(0) globules share protein and polysaccharide signatures, including lipooligosaccharides. Together these lines of evidence indicate that *Cba. tepidum*’s OMVs are one component of sulfur transport between cells and extracellular sulfur globules.

**IMPORTANCE:** All living cells must exchange material with their environment while maintaining cellular integrity. This is a particular challenge for materials that are not water soluble, yet many bacteria utilize insoluble materials for energy conservation and as nutrients for growth. Here we show that *Cba. tepidum* makes outer membrane vesicles and that these vesicles are likely involved in the exchange of material with extracellular elemental sulfur globules formed and consumed by *Cba. tepidum* as part of its energy metabolism based on oxidizing reduced sulfur compounds like hydrogen sulfide. These data expand our basic understanding of *Cba. tepidum*’s metabolism. As elemental sulfur is an industrial by-product with a limited number of uses, the information here may help enable the use of additional sulfur compounds by *Cba. tepidum* to drive the synthesis of biomass and/or specialty biochemicals from waste elemental sulfur by this autotrophic bacterium.

## INTRODUCTION

Many microbes interact with solids as part of their energy metabolism or to obtain elements required for growth. In either case, insoluble materials must pass through the cell envelope while maintaining cellular envelope integrity. Microbe-mineral interactions often occur in the context of redox reactions for energy metabolism. Examples include the production of extracellular sulfur globules (S(0)) by sulfide oxidation and their further oxidation to sulfate by phototrophic bacteria (1) or the utilization of Fe- and Mn-oxides as terminal electron acceptors (2, 3). These microbe-mineral interactions drive many biogeochemical cycles and the bioremediation of organic and inorganic pollutants (4). In Gram-negative bacteria, outer membrane vesicles (OMVs) are one mechanism used to interact with surfaces for multiple functional outcomes (5). For example, *Acinetobacter baumannii* OMVs enriched in TonB-dependent transporters enabled the acquisition of insoluble nutrients such as iron (6). Recently, OMVs containing a truncated form of lipopolysaccharide (LPS) were implicated as a key mechanism of *Geobacter sulfurreducens*’ elimination of uranium particles (7). Although OMVs have been intensively studied in heterotrophic Pseudomonadota, less is known about the production, composition, and physiological role of OMVs outside of this phylum.

OMVs are small (20-300 nm) spherical structures produced by Gram-negative bacteria via OM vesiculation (21). OMVs contain phospholipids, LPS or lipooligosaccharides (LOS), and outer membrane proteins (OMPs). OMVs also contain cargo that may include nucleic acids (DNA and RNA), periplasmic components, housekeeping proteins, and enzymes (22–24). OMVs were once regarded as cell debris or microscopy artifacts. Nowadays, OMVs have been shown to participate in diverse biological processes, such as transferring and delivering their components to the extracellular environment (25) and host-microbe interactions (26).

*Chlorobaculum tepidum (Cba. tepidum)* is an anaerobic photoautotrophic Gram-negative bacterium, but little is known about the composition and metabolism of the outer membrane (OM) in this organism or the Chlorobiaceae more broadly. *Cba. tepidum* is a model organism of the Green Sulfur Bacteria (GSB) due to rapid growth in culture (8), a sequenced and annotated genome (9), and genetic tools (10–12). *Cba. tepidum* utilizes a variety of inorganic sulfur compounds as electron donors to fix CO_2_ through the reverse TCA cycle (13, 14). *Cba. tepidum* both forms and consumes extracellular S(0) globules, but the details of how this occurs are unclear (1). However, the enzymes catalyzing S(0) production and consumption are believed to reside in the periplasm between the OM and cell membrane (15, 16). As S(0) is extracellular, there must be a transport mechanism for S(0) to cross the outer membrane during its production and consumption. Time-lapse microscopy of *Cba. tepidum* documents S(0) globules appearing in cultures with no visible connection to cells and increasing in size during growth on sulfide. S(0) globules produced by the wild type *Cba. tepidum* strain are roughly spherical and 1-2 μm in diameter (17, 20), while those produced by a mutant strain that cannot oxidize S(0) are larger, up to ∼8 μm in diameter (1). *Cba. tepidum* cells are typically ∼0.5 μm wide by 1-2 μm long (1, 17, 20).

*Cba. tepidum* grows well with elemental sulfur that it has synthesized, i.e. biogenic S(0) globules, as a sole electron donor and a sub-population of cells is tighly attached to S(0) during its utilization (1). *Cba. tepidum* S(0) globules have a proteome (1, 19) that includes proteins CT1320.1 and CT1305, OM proteins only found in the Chlorobiaceae and *Geobacter* spp., which also rely on interactions with insoluble minerals for energy metabolism (18). Biogenic S(0) globule utilization cannot be catalyzed by small metabolites or secreted soluble proteins as separating *Cba. tepidum* cells from biogenic S(0) with dialysis membranes ranging from 50 to 100 kDa prevented growth. Thus, relatively large proteins or complexes are required for biogenic S(0) utilization (1). Separately, time-lapse phase contrast microscopy showed that while some cells directly contact biogenic S(0), individual S(0) globules can be formed and consumed in cultures without direct contact by cells (17). Growth rates of cells that were not attached to S(0) globules were the same as those attached to S(0). Thus, material is released that allows cells at a distance from S(0) globules to grow as well as those that are attached. Spectroscopic characterization of *Cba. tepidum* S(0) globules indicated the presence of proteins and polysaccharides as an organic coating that was proposed to slow the conversion of amorphous biogenic S(0) to crystalline sulfur (20). *Cba. tepidum* biogenic S(0) globules stained with the membrane specific dye FM 1-43FX (20), suggesting that they contain lipids. Collectively, these observations from prior experiments suggest that *Cba. tepidum* S(0) globule formation and consumption involves components that contain outer membrane proteins, polysaccharides, lipids, in complexes larger than 100 kDa, but smaller than the resolution limits of phase-contrast microscopy. OMVs are a good candidate for these components.

Microbial production of extracellular S(0) associated with organic materials is well known, but mechanistic details underlying its formation are scarce (1, 17, 20, 27–29). S(0) transport by cytoplasmic membrane vesicles has been observed in Archaea, e.g. Gorlas *et al*. (30) reported that *Thermococcus* species produced vesicles containing S(0) solely when S(0) was added to the growth medium. They hypothesized this was to remove excess S(0) not used for growth, preventing the accumulation of toxic levels in the cytoplasm. Alternatively, the formation of organic associated amorphous S(0) has been proposed to be a precipitation process during sulfide oxidation in the presence of organics that can occur in spent medium in the absence of cells; a process called organomineralization (27, 28, 31, 32).

We hypothesize that *Cba. tepidum* produces OMVs that participate in S(0) transport between cells and extracellular S(0) globules. If the hypothesis is correct, then we predict proteomic analysis of OMV cargo should detect previously identified S(0) globule proteins (CT1305 and CT1320.1) and that there should be substantial overlap between OMV and S(0) globule proteomes. We would also expect lipid A in the form of lipopolysaccharide (LPS) or lipooligosaccharide (LOS) should be present in both OMVs and S(0) globules. Finally, we would also expect that OMV abundance will be higher when S(0) is actively being produced or consumed by *Cba. tepidum*. Here, we test these predictions through a combination of OMV purification, proteomic and LPS analysis of both OMVs and S(0) globules, and OMV quantification under different growth conditions. Raman spectroscopy was used to investigate the chemical signatures of OMVs and S(0). The data are consistent with a model where OMVs participate in extracellular S(0) globule formation characteristic of the Chlorobiota. Furthermore, the results reported here provide significant new information on OM structure and function outside the Pseudomonadota. In the following sections, we use the term ‘particle’ to describe materials that were analyzed to determine if they were OMVs or not. We only use the term OMV after presenting data that their size and molecular composition of *Cba. tepidum* extracellular particles is consistent with them indeed being OMVs.

## RESULTS AND DISCUSSION

### *Cba. tepidum* secretes extracellular particles during growth

While OMV secretion is considered a universal process among Gram-negative bacterial species (33, 34), most observations to date are from heterotrophic members of the phylum Pseudomonadota, formerly Proteobacteria (35–37). *Cba. tepidum* is a member of the phylum Chlorobiota, which is sister to the phylum Bacteroidota and distantly related to the Pseudomonadota. Therefore, we examined *Cba. tepidum* cultures for extracellular particles by microfluidic resistive pulse sensing (MRPS) analysis of 0.2 μm culture filtrates after 8 h (lag phase), 14 h (early-exponential phase), and 24 h (late-exponential phase) of growth. Briefly, MRPS detects and measures particles based on changes in electrical resistance as particles pass through a microfluidic pore that is charged held at a set voltage. The size range of particles detectable in these experiments was from ∼40 nm to 400 nm. The number of particles detected was highest during the early-exponential growth phase (14 h of growth, Fig. 1A) and these particles contained protein (Fig. 1B). Protein and particle concentrations displayed the same pattern across growth phases. We chose early-exponential phase samples to produce large amounts of particles for more detailed analysis because of the larger particle concentration and to minimize capturing debris deriving from cell death or nutrient limitation during later growth stages (41–43). Direct observation of *Cba. tepidum* cells in 14 h culture samples by TEM found vesicles at the outer membrane (Fig. 2A, B) suggesting that *Cba. tepidum* actively produces particles via OM blebbing. Particles purified from the early-exponential imaged by negative stain TEM were spherical and displayed a cavity surrounded by a stained membrane (Fig. 2C, D). These data demonstrate particle production by *Cba. tepidum* shares similarities with OMV production: growth phase-dependence, particles appear to be produced by OM blebbing, and TEM images resembling a deflated ball.

**Figure 1.**
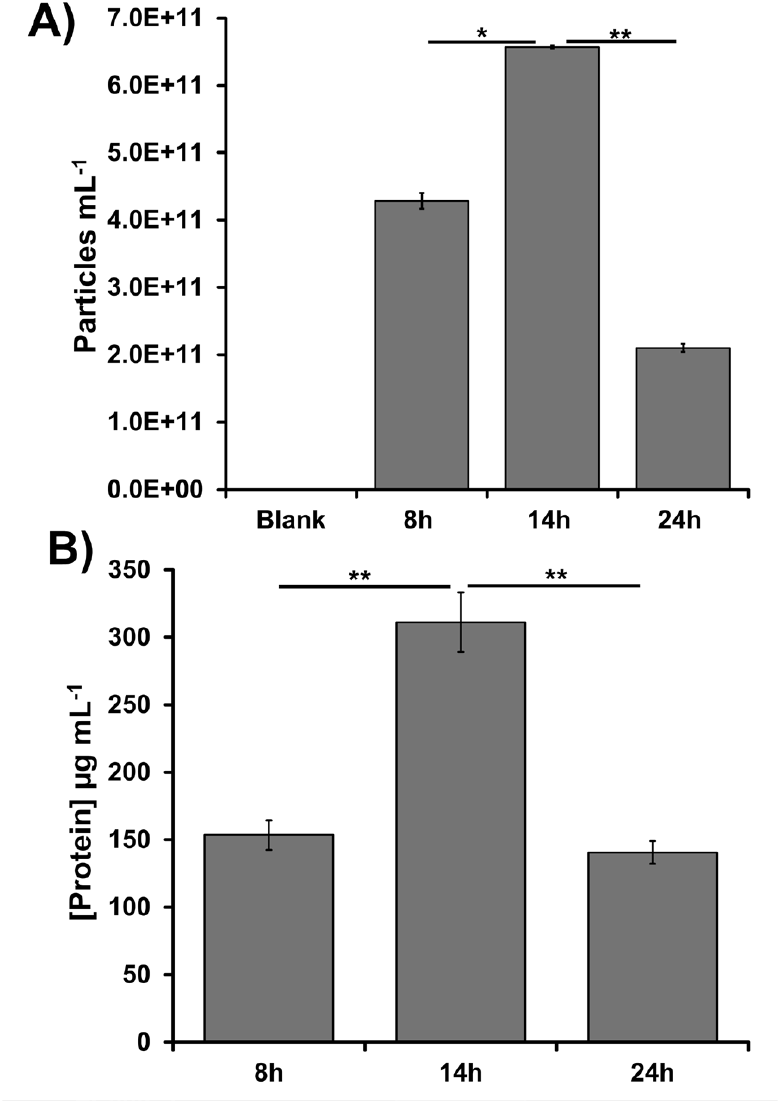
The effect of *Cba. tepidum* growth phase on extracellular particle concentration in cultures and protein concentration in purified particle preparations. See Fig. S1A for details on purification. **A)** Extracellular particles concentration from different growth phases. **B)** Total protein concentration of particles purified at different growth phases. Error bars represent the standard error, n = 3 with * indication p < 0.05 and ** indicating p < 0.01 (t-test, two tailed, unequal variance).

**Figure 2.**
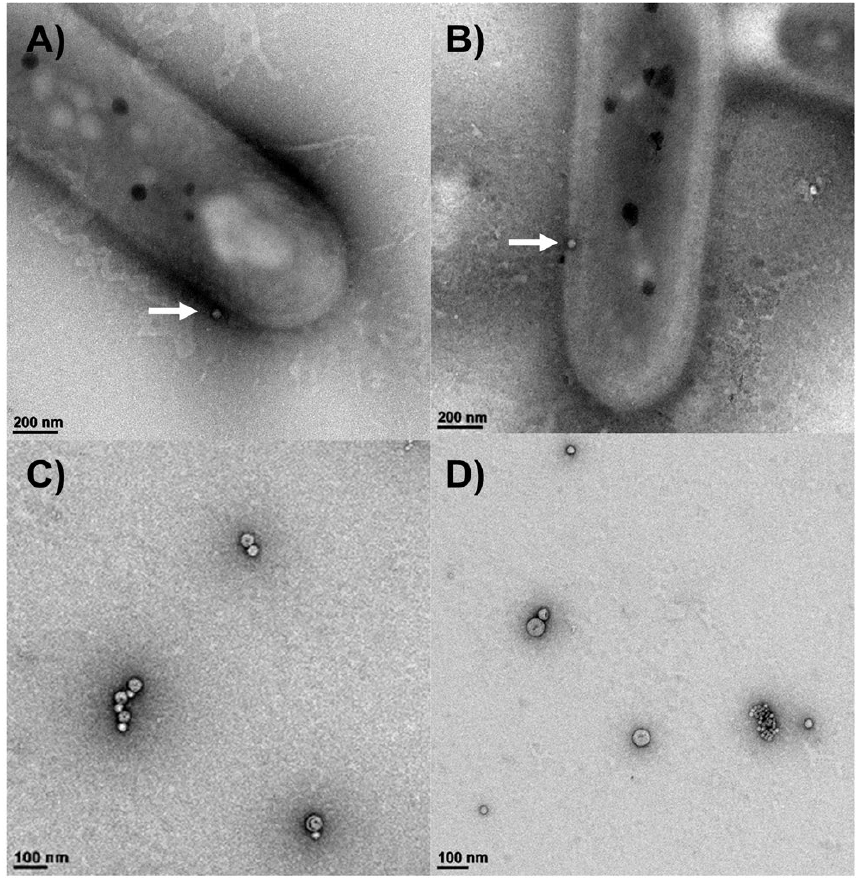
Transmission electron microscopy (TEM) images of *Cba. tepidum*’s OMVs. **A)** and **B)** TEM images show spherical OMVs budding out of rod-shaped *Cba. tepidum* after 8h and 14h of growth, respectively (white arrows). **C)** and **D)** TEM analysis of OMVs purified via serial ultracentrifugation from 14 h old *Cba. tepidum* culture.

### Particles produced by *Cba. tepidum* are the same size as OMVs and contain lipooligosaccharides

*Cba. tepidum* can grow on sulfide, thiosulfate, or biogenic S(0) as electron donors. Therefore, we examined whether particles differed when they were purified (Fig. S1) from cultures grown on single electron donors or the combination of sulfide and thiosulfate in Pf-7, the standard growth condition (1, 8). *Cba. tepidum* growth rate was not significantly different between the growth conditions (data not shown). Particles were characterized both by transmission electron microscopy (TEM) and MRPS. TEM imaging requires sample concentration by ultracentrifugation and chemical treatment prior to imaging, while MRPS provides information on particle size and concentration after 0.2 µm filtration of the supernatant as the sole sample preparation step (Fig. S1B). Therefore, we utilized MRPS to compare the concentration and size of particles across growth conditions, while TEM image analysis was used to check particle morphology and size. Mature S(0) globules produced by *Cba. tepidum* are larger than 0.2 μm and should not be present in these samples (1, 17). The particle size distributions by TEM (10-105 nm) and MRPS (40-125 nm) overlapped (Fig. S2), with the difference likely being the size limit of detection for the MRPS cartridge being ∼40 nm. Particles produced when thiosulfate was the sole electron donor were slightly larger by MRPS than particles released under other conditions, but the size distribution of *Cba. tepidum* particles purified from all conditions overlapped and is consistent with published values for OMVs (Fig. 3A). Regardless of the electron donor, the counted particles contained protein, with a linear relationship between particle and protein concentrations (Fig. 3B), suggesting that particles have a constant concentration of protein per unit of particle volume

**Figure 3.**
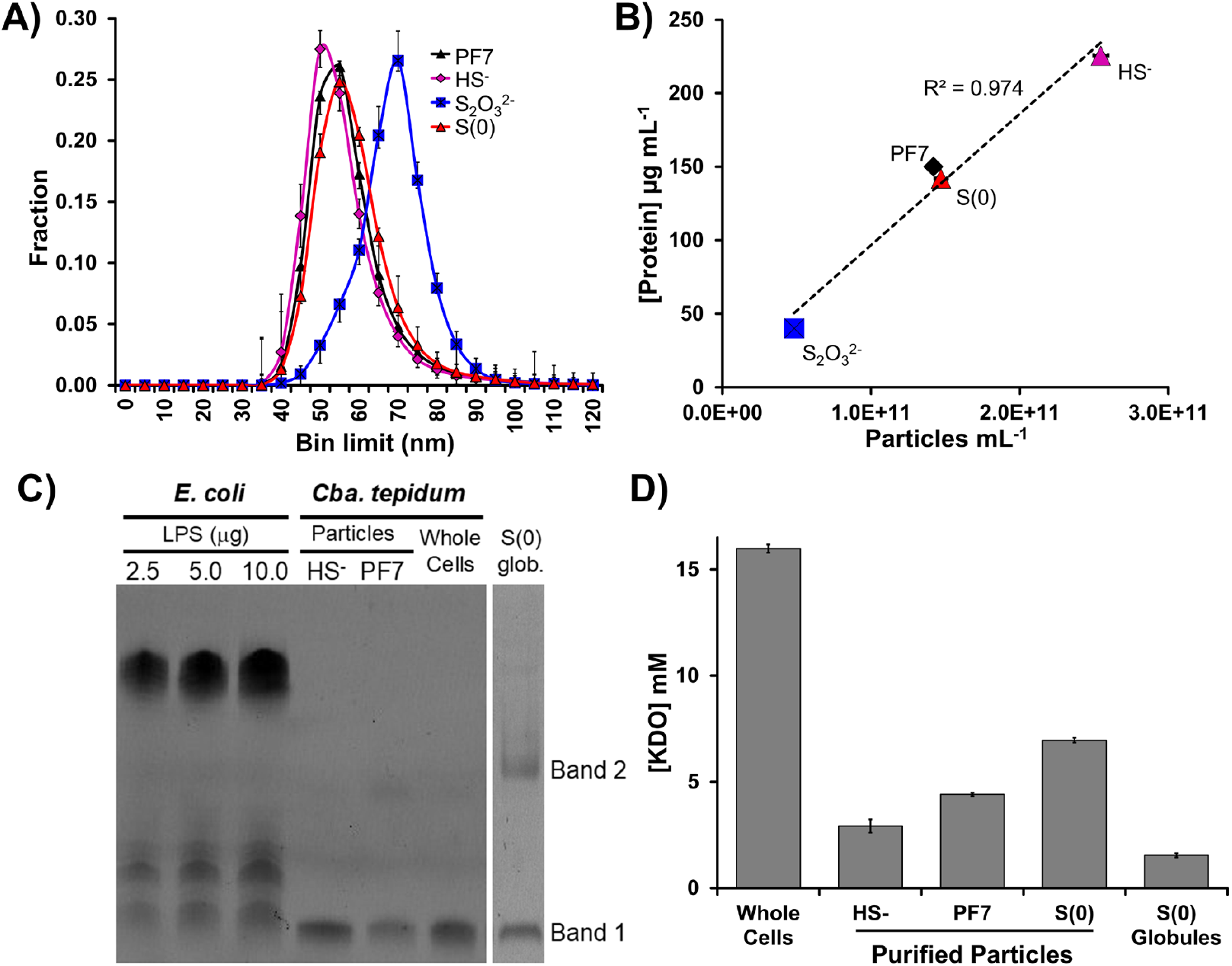
Analysis of *Cba. tepidum* extracellular particles for hallmark characteristics of OMVs including size (A), protein (B), LPS or LOS (C), KDO (D).

Lipopolysaccharide (LPS), the major component of the outer leaflet of the OM, plays a crucial role in the stability and permeability of the OM and should be present in *Cba. tepidum* particles if they are OMVs. Therefore, we tested whether particles and biogenic S(0) produced by *Cba. tepidum* contain LPS. The *Cba. tepidum* genome does contain genes for lipid A biosynthesis (data not shown), but the structure of the outer membrane lipids has not been established to our knowledge. Tricine SDS-PAGE was used to visualize material produced by an LPS purification protocol: material from *Cba. tepidum* whole cells or purified particles (Fig. S1A) resolved in a single low molecular weight band (Fig. 3C) while material purified from biogenic S(0) resolved into two bands (Fig. 3D), with the lower molecular weight band migrating identically to that from cells and particles. The *E. coli* purified LPS standard produced a laddered pattern characteristic of an O antigen (Fig. 3C). Therefore, we conclude that *Cba. tepidum* makes lipooligosaccharide (LOS), composed of lipid A and an inner core oligosaccharide (band 1). *Bacteroides th*e*taiotamicron*, a member of the phylum Bacteroidota more closely related to *Cba. tepidum* than *E. coli*, also synthesizes LOS (44). Similarly, Meißner *et al*. (45) showed that purified OM lipid from *Chlorobium vibrioforme* had no O antigen repeating units, i.e. it is LOS. The identity of band 2 observed in biogenic S(0) is unclear, but the staining method used is specific for sugars, e.g. glycolipids like those in LPS and LOS. These samples did not contain any material that stained with SYPRO Ruby (data not shown), indicating that the LPS preparations were protein-free.

The identity of the material in the LPS preparations was further established by mass spectrometry and an HPLC assay for the inner core sugar 2-ketodeoxy-octulosonic acid (KDO). Masses consistent with lipid A were observed in electrospray ionization Fourier transform MS after mild acid hydrolysis of LOS isolated from whole cells, OMVs, and biogenic S(0). Different forms of lipid A were detected in each sample. The hepta-acyl form was found in WC, HS-OMV, and S(0) (*m/z* 1851, 1894, 1925, 1926, 1912), while the penta-acyl form was found only in Pf7-OMV (*m/z* 1659, 1705, 1795). The observed diversity of lipid A forms in *Cba. tepidum* suggests some differences in their acyl chain length and number depending on growth condition, which may in turn affect or be a result of OMV production (46–48). KDO was detected in all samples tested, with the highest level found in whole cells (Fig. 2E). No KDO was detected in any mock samples: PBS and chemical sulfur purchased from Sigma-Aldrich using the same reagents and buffers for preparing the samples. Taken together with the data above on particle size, TEM appearance, and protein content, the detection of glycolipid staining, Lipid A, and KDO in particles produced by *Cba. tepidum* confirms that these are indeed OMVs and this term will be used hereafter. Detection of these signatures in biogenic S(0) supports a hypothesis where OMVs may be one route of transfer of S(0) to extracellular S(0) globules, which is consistent with prior data on the composition of *Cba. tepidum* S(0) globules (1, 19, 20).

### Proteome analysis of *Cba. tepidum* OMVs versus whole cells and biogenic Sulfur

Previously, we showed that *Cba. tepidum* S(0) globules contained the proteins CT1305 and CT1320.1 (1), which are only shared with *Geobacter* spp. that also interact with insoluble phases for energy metabolism. Deeper analysis of these extracts by shotgun proteomics indicated that biogenic S(0) contained a significant proteome of at least 96 proteins (Supplemental Table S1), which is lower than the 696 S(0)-associated proteins recently identified by Lyratzakis *et al*. (19). Here, we generated new shotgun proteomic datasets for *Cba. tepidum* OMVs, biogenic S(0), and whole cells of *Cba. tepidum* to control for S(0)-attached cells that are difficult to separate from biogenic S(0) (1). The whole cell samples were diluted to an equal concentration of bacteriochlorophyll (Bchl) *c* observed in biogenic S(0). This assumes that Bchl *c* in biogenic S(0) is from adherent cells and was done to normalize the total amount of protein in the whole cell samples to that of adhered cells in biogenic S(0). This is needed for label-free quantification comparisons between samples. A total of 1,447 unique proteins were identified across all samples (Fig. 4A), with 757 of these detected in S(0) globules, somewhat more than the number observed by Lyratzakis *et al*. (19). OMVs from thiosulfate-only cultures only contained two proteins, CT0068 and CT2144, which are both outer membrane beta-barrel proteins, suggesting that electron donors might play a key role in shaping the cargo carried by OMVs. For example, Liu *et al*. (49) showed that differences in redox potential might impact the packaging of specific molecules (lipids, enzymes, proteins, toxins) into OMVs. Based on the limited proteome in thiosulfate-OMVs, this condition was excluded from further comparative analysis.

**Figure 4.**
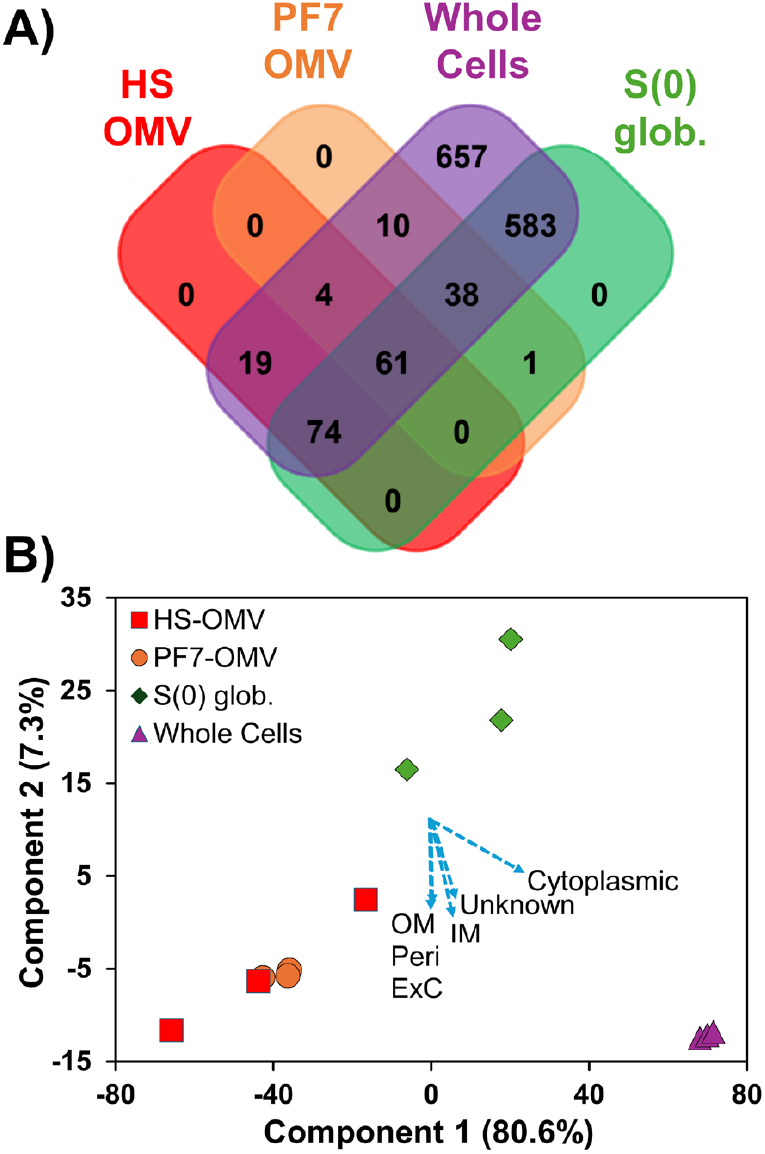
Comparative proteome analysis of *Cba. tepidum* whole cells, OMVs from cultures grown on sulfide (HS-OMV) or sulfide + thiosulfate (PF7-OMV), and biogenic sulfur globules (S(0) glob.). **A)** Venn diagram indicating the distribution of 1,447 proteins across sample group (n = 3 samples per group). **B)** Principal component analysis score plot of samples based on label free quantification. The first and the second principal components (PC) account for 80.6% and 7.3% of the variability in the data, respectively. Loadings for PC1 and PC2 are displayed (blue dashed arrows) for proteins grouped by PSORTb predicted localization: Cytoplasmic, Unknown, Inner Membrane (IM), Outer Membrane (OM), Periplasmic (Peri), and Extracellular (ExC).

OMV proteomes from cultures grown on sulfide (HS-OMVs), sulfide + thiosulfate (PF7-OMVs), purified biogenic S(0) globules, and diluted whole-cells shared 31 proteins detected in at least two independent samples for each group (Table 1), which we considered to be the OMV core proteome. This is a more conservative estimate than the Venn diagram (Fig. 4A), which only required detection of a protein in one of the three independent samples within a group. The functional annotations of the 31 proteins include porin activity, electron transfer activity, and trafficking of unfolded proteins. Furthermore, two periplasmic proteins, SurA (CT2264) and DegP (CT1447), identified in the initial S(0) globule proteome (Supplemental Table S1), were also found in HS-OMVs. Based on experimental evidence (50) and a computational model (51), SurA is believed to be involved in the trafficking of unfolded outer membrane proteins, while DegP has protease activity and is involved in the degradation of misfolded proteins, preventing the accumulation of waste in the periplasm. CT1305, one of two poorly characterized proteins shared with *Geobacter* spp. and detected in the initial S(0) proteome (Supplemental Table S1), was detected in PF7-OMVs, S(0) globules, and whole cells, but not HS-OMVs, so it was not included in the OMV core. The other, CT1320.1, was not detected in any of the samples.

**Table 1.**
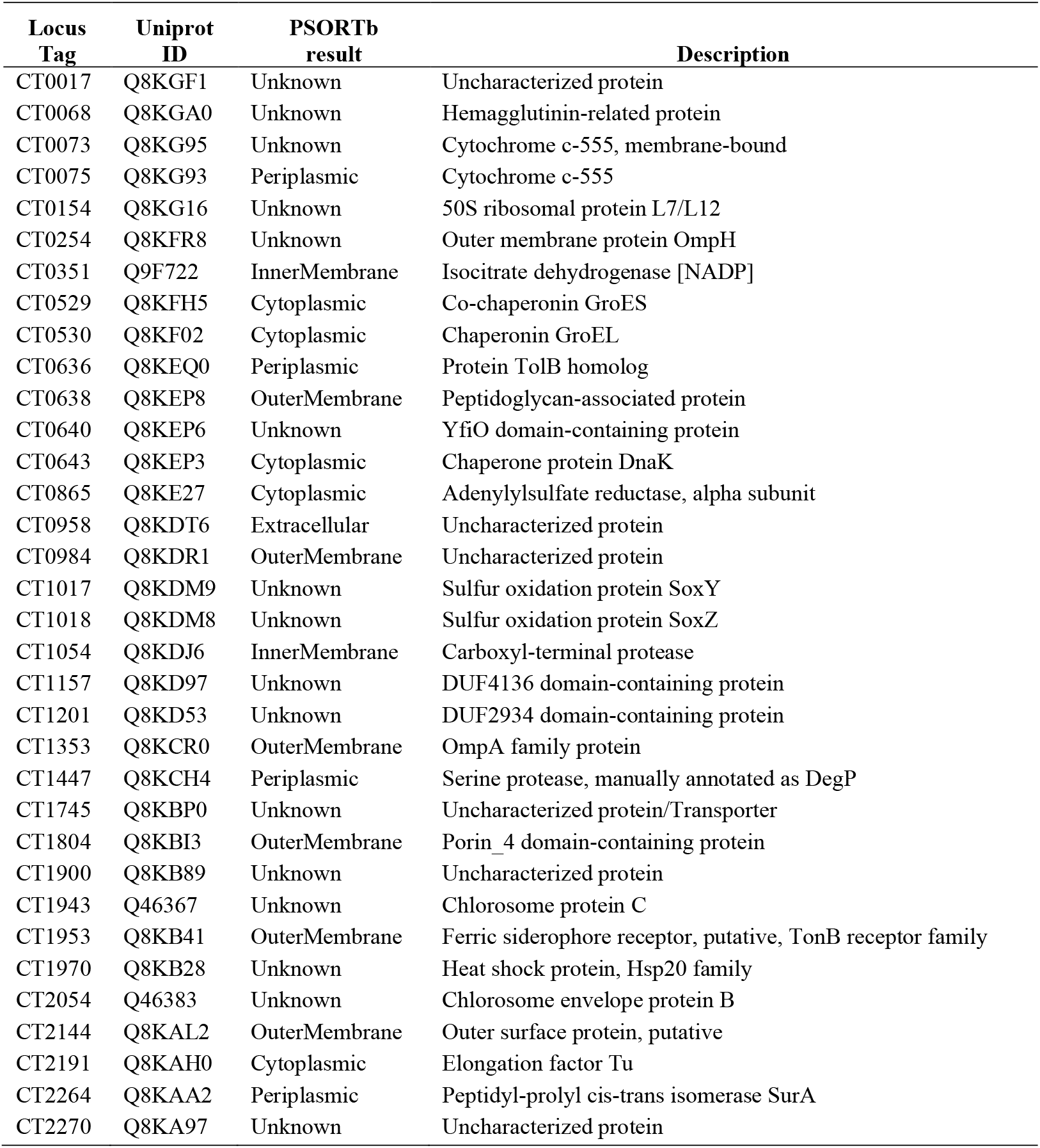
OMV core proteins identified in ≥ 2 independent samples from each group (whole cells, S(0) globules, HS-OMV, and PF7-OMV) by label-free quantitative proteomics.

| Locus Tag | Uniprot ID | PSORTb result | Description |
| --- | --- | --- | --- |
| CT0017 | Q8KGF1 | Unknown | Uncharacterized protein |
| CT0068 | Q8KGA0 | Unknown | Hemagglutinin-related protein |
| CT0073 | Q8KG95 | Unknown | Cytochrome c-555, membrane-bound |
| CT0075 | Q8KG93 | Periplasmic | Cytochrome c-555 |
| CT0154 | Q8KG16 | Unknown | 50S ribosomal protein L7/L12 |
| CT0254 | Q8KFR8 | Unknown | Outer membrane protein OmpH |
| CT0351 | Q9F722 | InnerMembrane | Isocitrate dehydrogenase [NADP] |
| CT0529 | Q8KFH5 | Cytoplasmic | Co-chaperonin GroES |
| CT0530 | Q8KF02 | Cytoplasmic | Chaperonin GroEL |
| CT0636 | Q8KEQ0 | Periplasmic | Protein TolB homolog |
| CT0638 | Q8KEP8 | OuterMembrane | Peptidoglycan-associated protein |
| CT0640 | Q8KEP6 | Unknown | YfiO domain-containing protein |
| CT0643 | Q8KEP3 | Cytoplasmic | Chaperone protein DnaK |
| CT0865 | Q8KE27 | Cytoplasmic | Adenylylsulfate reductase, alpha subunit |
| CT0958 | Q8KDT6 | Extracellular | Uncharacterized protein |
| CT0984 | Q8KDR1 | OuterMembrane | Uncharacterized protein |
| CT1017 | Q8KDM9 | Unknown | Sulfur oxidation protein SoxY |
| CT1018 | Q8KDM8 | Unknown | Sulfur oxidation protein SoxZ |
| CT1054 | Q8KDJ6 | InnerMembrane | Carboxyl-terminal protease |
| CT1157 | Q8KD97 | Unknown | DUF4136 domain-containing protein |
| CT1201 | Q8KD53 | Unknown | DUF2934 domain-containing protein |
| CT1353 | Q8KCR0 | OuterMembrane | OmpA family protein |
| CT1447 | Q8KCH4 | Periplasmic | Serine protease, manually annotated as DegP |
| CT1745 | Q8KBP0 | Unknown | Uncharacterized protein/Transporter |
| CT1804 | Q8KBI3 | OuterMembrane | Porin_4 domain-containing protein |
| CT1900 | Q8KB89 | Unknown | Uncharacterized protein |
| CT1943 | Q46367 | Unknown | Chlorosome protein C |
| CT1953 | Q8KB41 | OuterMembrane | Ferric siderophore receptor, putative, TonB receptor family |
| CT1970 | Q8KB28 | Unknown | Heat shock protein, Hsp20 family |
| CT2054 | Q46383 | Unknown | Chlorosome envelope protein B |
| CT2144 | Q8KAL2 | OuterMembrane | Outer surface protein, putative |
| CT2191 | Q8KAH0 | Cytoplasmic | Elongation factor Tu |
| CT2264 | Q8KAA2 | Periplasmic | Peptidyl-prolyl cis-trans isomerase SurA |
| CT2270 | Q8KA97 | Unknown | Uncharacterized protein |

We assessed the relationship between protein composition and sample characteristics using Principal Components Analysis (PCA, Fig. 4B) of log2 transformed label-free quantification abundance data. Our analysis showed that the first two components accounted for a high proportion of the total variation (87.9%) between samples, with those of each group clustering together. The OMV and S(0) samples were well separated from whole cells on the PC1 axis (80.6% of variation). The OMV and S(0) samples also separated along the PC2 axis (7.3% of variation), indicating that they are distinct from one another. Proteins were grouped by their predicted subcellular localization and the cumulative loadings for these groups (Fig. 4B, blue dashed arrows) indicate that cytoplasmic proteins are responsible for separation of the whole cell samples from the others. The loadings for predicted outer membrane, periplasmic and extracellular proteins are very similar and aligned along the PC2 axis, suggesting that they strongly contribute to separation of samples in this dimension.

### The effect of electron donors on OMV abundance

The relationship between the proteomes of OMV and biogenic S(0) prompted us to examine whether sulfide concentration modulated OMV production. We hypothesized that increasing sulfide concentrations will cause *Cba. tepidum* to release more vesicles to prevent accumulation of S(0) produced by sulfide oxidation, thus playing a key role in sulfur detoxification as proposed for Archaea (30). The presence of the SurA and DegP may indicate that S(0) induces unfolded protein stress as another signal for OMV production. *Cba. tepidum* was grown on a range of sulfide concentrations (2-7 mM), and OMVs from each sulfide concentration were analyzed by MRPS. The data showed that, as sulfide concentrations are increased, particle concentration increases versus 2 mM sulfide (Fig. 5A) by 2.8-fold at 4 mM sulfide and ∼10.7-fold at 5 mM, with no further significant increase at 7 mM. This result suggests that growing *Cba. tepidum* at 5 mM sulfide produced the maximum number of particles. MRPS also provides size information and documented an increase in median particle diameter from 61 nm at 2 mM sulfide to 65-67 nm at 4-5 mM while those from 7 mM cultures were 88 nm in diameter. OMV yields can be expressed in different terms, such as lipid concentration, amount of protein, total dry weight harvested, or counts by TEM, making the comparison of OMV yield using all these methods simultaneously difficult because the measurements also encompass non-vesicular materials (52, 53). TEM comparison of vesicles from 4 mM and 7 mM cultures independently confirmed the increase in OMV concentration in the 7 mM sulfide OMVs (Fig. 5 C, D). OMVs from 7 mM sulfide cultures were larger in the TEM images than those from 4 mM sulfide, but this was not significant (p = 0.12, t-test, 1-tailed, homoscedastic, n = 13 particles each). We feel the MRPS data strongly supports an increase in OMV size with increased sulfide due to the larger number of particles measured and a consistent increase of larger sizes at all sulfide concentrations in this experiment (Fig. 5B, points in the 80-110 nm range). The peak position for 7 mM may be influenced by clumping indicated by TEM (Fig. 7D), but clumping could also be a TEM sample preparation artefact. Finally, we directly compared OMVs from these samples at higher magnifications (Fig. S3) and are confident that the OMVs have the same overall structure across all samples. Collectively, these data confirm that OMV numbers and sizes vary with increased sulfide concentrations.

**Figure 5.**
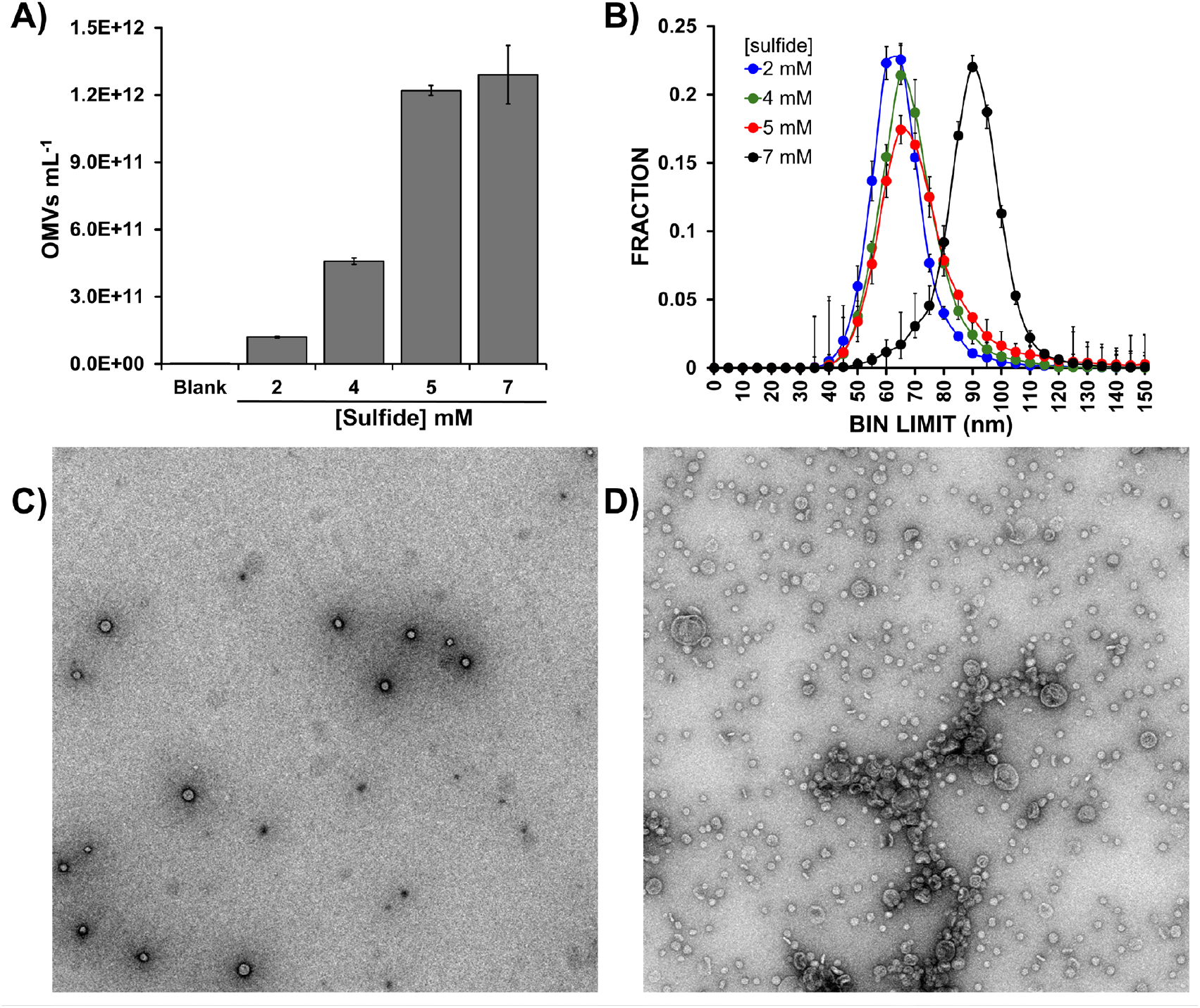
OMVs concentration and size vary with increased sulfide (HS) concentration. A) OMV concentrations measured after 14 h of growth with different concentrations of sulfide as the sole electron donor. B) MRSP-based size distribution of particles purified from the cultures in panel A (see Fig. S1A for purification details). C) and D) TEM analysis of OMVs purified from 4 mM and 7 mM sulfide cultures in panel A corresponding to the green and black traces in panel B.

**Figure 6.**
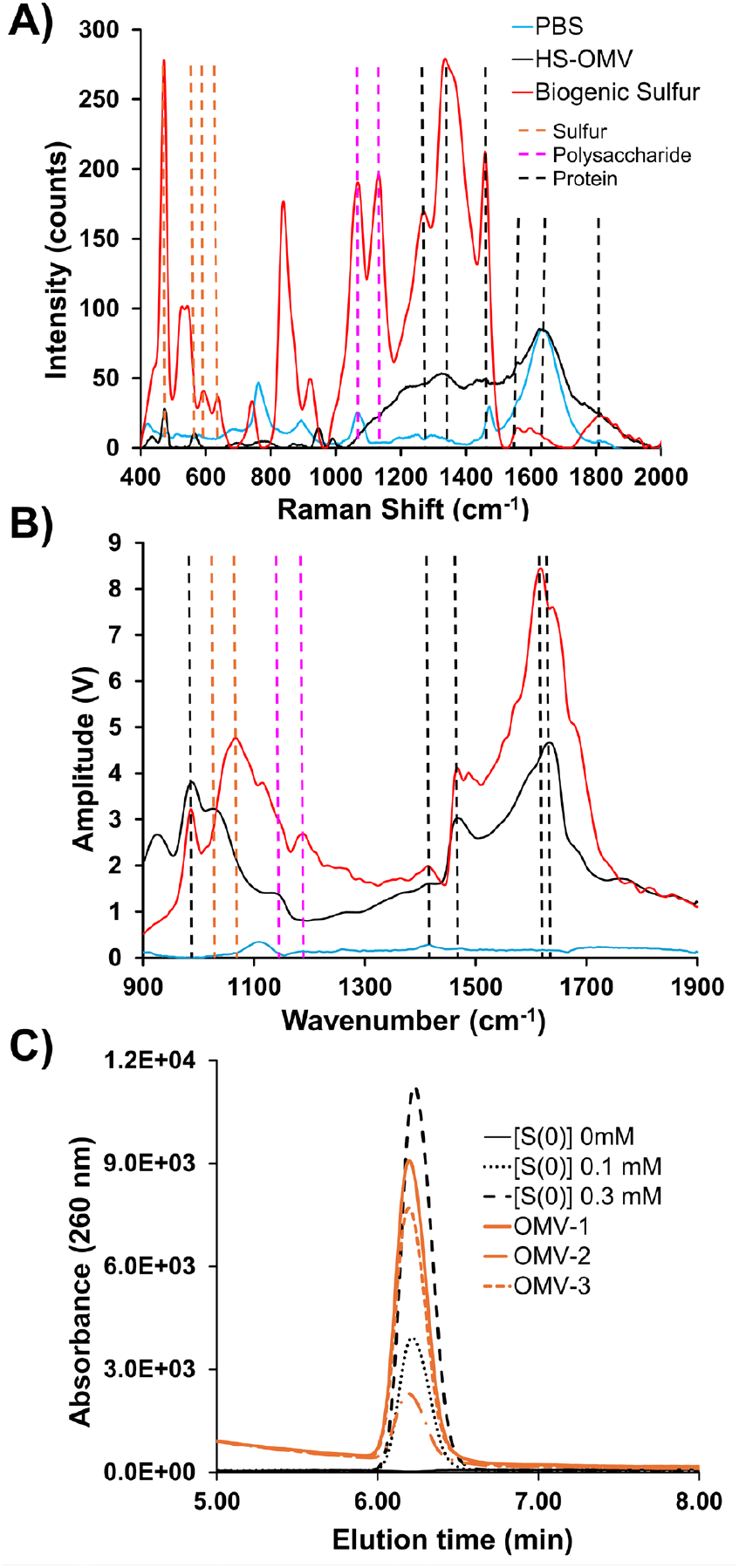
OMVs and S(0) share multiple characteristic. Characteristic Raman scattering **(A)** or IR absorption **(B)** spectra for purified OMVs (black), biogenic S(0) (red), and PBS (blue). Scattering (A) or absorption (B) wavenumbers are indicated by vertical dashed lines for sulfur (orange), polysaccharide (pink), and protein (black). **(C)** Chromatograms of elemental sulfur standards (0-0.3 mM S(0), black traces) and OMV’s purified from three independent 7 mM sulfide *Cba. tepidum* cultures (orange traces). The standards indicate the peak from OMV samples co-migrates with authentic S(0) and the replicates are within the range of detection.

**Figure 7.**
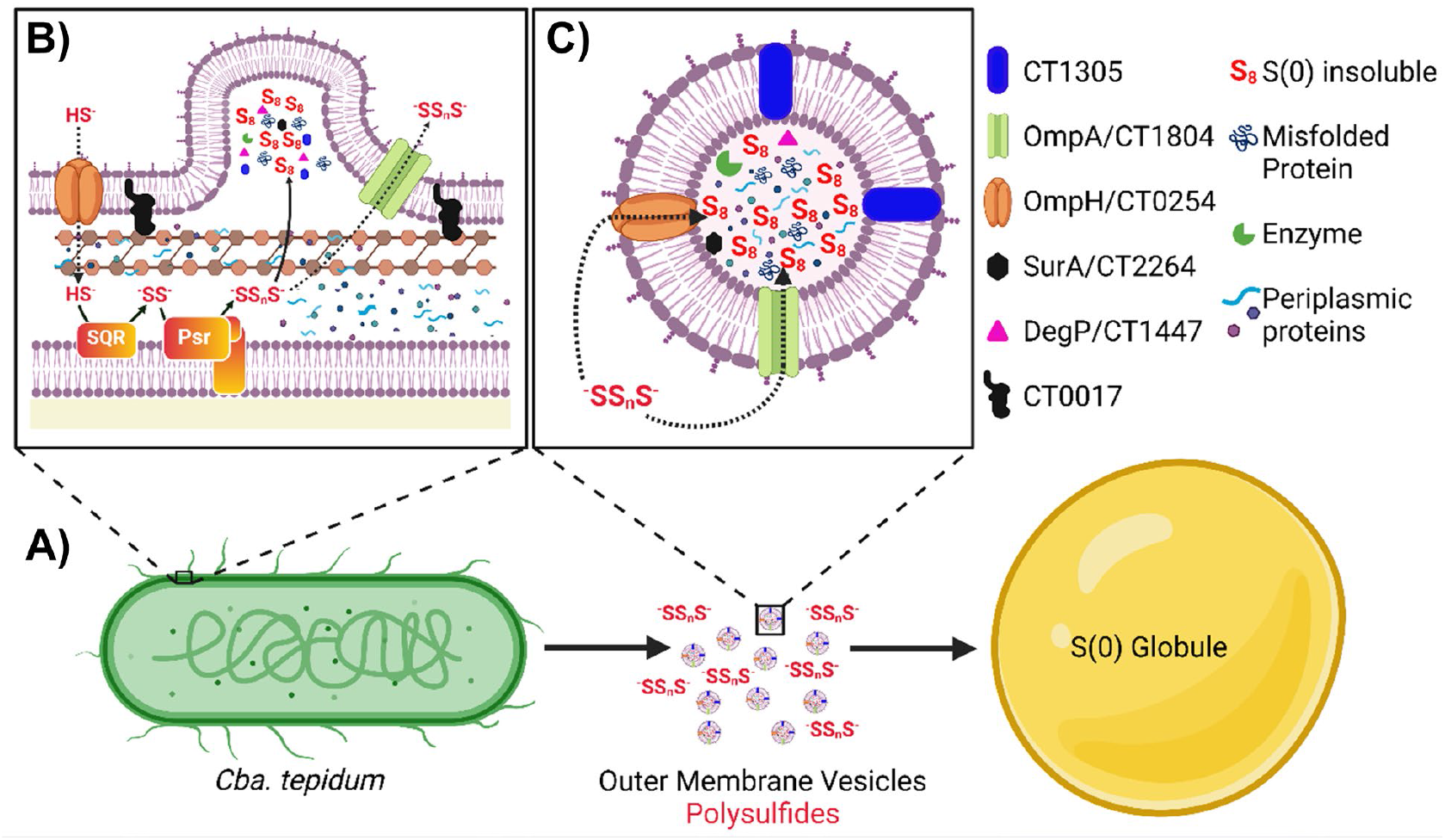
Summary of the proposed OMV S(0) transport mechanism. A) *Cba. tepidum* produces both OMVs and polysulfides during growth that contribute to the growth of extracellular S(0) globules. B) Enlarged view of the periplasm and OM with increased detail. Sulfide oxidation may induce unfolded protein stress that leads to OMV production. The dashed arrows for sulfide and polysulfide indicate possible routes to enter and exit either OMVs or S(0) globules. SQR = sulfide:quinone oxidoreductase (CT0117 and CT1087), Psr = putative polysulfide oxidoreductase complex (CT0496-CT0494). C) Enlarged view of a *Cba. tepidum* OMV with selected cargo molecules detected by proteomics analyses denoted at the right. Created in BioRender https://BioRender.com/bzod9yf

### Molecular Composition of S(0) globules and OMVs

The molecular composition of S(0) globules or OMVs was probed with Raman microscopy and atomic force microscopy coupled with infrared (AFM-IR) spectroscopy. Raman microscopy is a non-destructive method that gives a linear correlation of compound concentration to signal strength (31). Raman microscopic analysis of biogenic S(0) and HS-OMV samples deposited on aluminum revealed signals at 425-550 cm^-1^ (S-S, Sulfur), 580-680 cm^-1^ (C=S, thiocarbonyl), 1050-1130 cm^-1^ (carbohydrate), 1220-1290 cm^-1^ (amide III), and 1660-1680 cm^-1^ (amide I) (Fig. 6A). Different intensities of characteristic spectral regions showed that Sulfur, polysaccharides, and protein concentration were higher in S(0) globules than in HS-OMVs. S(0) may be added to a growing S(0) globule by an alternative route, possibly by excreted polysulfides as suggested by Marnocha *et al*. (17). Porin proteins (e.g. OmpA and OmpH) detected in *Cba. tepidum* OMVs may provide channels through which anionic polysulfide can diffuse and increase S(0) concentration. Additional organic materials may also be accumulating in S(0) globules by organomineralization processes (32).

AFM-IR spectra were collected for individual S(0) globules and OMVs. These show characteristic bands for proteins (1400-1650 cm^-1^), polysaccharides (1150-1200 cm^-1^), and sulfoxide (1030-1060 cm^-1^) (Fig. 6b). Sulfur is an essential component of proteins; thus, sulfoxide found in these samples could originate from sulfur-containing amino acids such as methionine, cysteine, and homocysteine. However, Steudel *et al*. (54) collated multiple lines of evidence to indicate that bacterial S(0) globules contain long-chain polythionates, which may also be the source of this signature. The bands around 1650 and 1460 cm^-1^ correspond to amide I and II vibrations of the peptide backbone, respectively. The presence of polysaccharides and proteins is confirmed by the presence of the aliphatic group C=O bonds (around 1660 cm^-1^) (55), the weak peak around 1330-1430 cm^-1^ characteristic for a hydroxyl group, and a weak C-H peak at 1370-1390 cm^-1^. The pyranose band was observed in the regions around 1150 and 1192 cm^-1^. Though there are advantages to the usage of AFM-IR for single S(0) and OMVs characterization, smaller vesicles or S(0) (15-50 nm) may not provide a strong IR signal for a consistent characterization. Furthermore, the range of IR wavelengths available on the AFM IR instrument was 950-1900 cm^-1^, which would miss the strongest absorbance peaks for S-S bonds that occur below 550 cm^-1^. The Raman instrument was capable of probing this region and detected the expected S-S scattering, increasing confidence in the interpretation of these data.

### OMVs from high sulfide cultures contain S(0)

While Raman and IR spectroscopy analyses were consistent with elemental sulfur being a component of HS-OMVs, the most basic question is whether purified OMVs contain elemental sulfur in significant amounts. Thus, HPLC of hexane extracts was used to quantify sulfur found in HS-OMVs. HPLC analysis confirmed the presence of elemental sulfur in HS-OMV (Fig. 6C) that were harvested from 20 mL cultures and extracted with 1 mL of hexanes. S(0) in the HS-OMV fraction was 94 ± 6 μM after accounting for the centrifugal concentration of the OMV sample before extraction.

We calculated the maximum possible [S(0)] in the OMV fraction of a *Cba. tepidum* culture with 7 mM sulfide as follows. The particle concentration of purified OMVs at the time of sampling (14 hrs) was 1.29 x 10^12^ particles/mL and the mean OMV diameter was 88 nm. The volume per OMV, assuming they are spherical, is 3.82 x 10^-16^ mL. The total OMV volume per mL of culture is then 4.92 x 10^-4^ mL. Different density values have been reported for biogenic S(0), 1.22 g cm^-3^ (56), 1.31 g cm^-3^ (57) or 2.08 g cm^-3^ (58). Based on these values, the predicted [S(0)] if OMVs were composed of solely of biogenic S(0) ranged from 18.8-32.0 mM, which is far more sulfide than was provided to the culture. The observed S(0) concentration in OMVs is ∼1-2% of the 7 mM sulfide provided to the culture, which is consistent with the hypothesis that OMVs are a transient pool of S(0) with the presence of a complex proteome and LPS in these samples confirming that S(0) is not the only cargo.

## Conclusions

The data above clearly indicate that *Cba. tepidum*, and by extension the Chlorobiaceae, synthesize LOS, secrete OMVs, and that OMVs likely contribute to material exchange between cells and S(0) globules during their production and consumption. In addition, the data presented here partially support our hypotheses that previously identified S(0) globule proteins would be found in OMVs and that there should be substantial overlap between OMV and S(0) globule proteomes. While CT1305 was found in OMVs, CT1320.1 was absent; such differences in OMV protein cargo might be related to the bacterial growth stage at which OMVs were isolated or difference in the proteomic sample preparation methods used. Several other pieces of evidence agree with our hypotheses and indicate the roles of OMVs in sulfur metabolism: (i) OMVs and S(0) globules share a substantial number of proteins; (ii) OMV abundance is higher when S(0) is produced and consumed by *Cba. tepidum*; and (iii) OMVs and S(0) share a significant number of chemical constituents, including elemental sulfur as documented by multiple independent methods.

Taking the observations made here and by others, we propose a mechanism where *Cba. tepidum* oxidizing sulfide produces OMVs to release accumulating S(0) and unfolded proteins from the periplasmic space (Fig. 7). Following the accumulation of these cargos, the OM can bulge outwards and pinch off as vesicles. We hypothesize that porins in the core OMV proteome (e.g. e.g. OmpA/CT1353 or OmpH/CT0254) in the growing S(0) globules may also allow for additional S(0) to accumulate perhaps by diffusion of polysulfides from cells. Clearly, additional mechanisms of transport and/or globule maturation occur, otherwise the chemical and proteome compositions of OMVs and S(0) globules would be nearly identical. Previous measurements of *Cba. tepidum* S(0) globule elemental composition indicate that they are > 99% S (1). A maturation process may explain why purified S(0) globules contain higher relative amounts of S, polysaccharides, and amides than OMVs based on the IR and Raman spectroscopy data presented above. A simple possibility could be the exclusion or expulsion of water and hydrophilic OMV cargos if multiple S(0) payloads condense by hydrophobic attraction. Experiments to observe OMV fusion and S(0) globule maturation in over time from purified OMVs and determining if S(0)-loaded OMVs can be used for growth by *Cba. tepidum* are obvious next steps to further dissect this process.

## MATERIALS AND METHODS

### Bacterial strain and growth conditions

*Cba. tepidum* strain WT2321 was used in all cultures and grown in Pfennig’s (Pf-7) medium, as previously described by (59) with an anaerobic headspace composed of 5% CO_2_ and 95% N_2_ passed through a heated copper scrubber. *Cba. tepidum* cultures used for OMVs isolation were incubated at 47°C with light provided by incandescent bulbs at an intensity of 20 µmol photons m^-2^ s^-1^ measured with a light meter equipped with a quantum PAR sensor (LI-COR, Lincoln, NE). *Cba. tepidum* liquid cultures were grown to different growth phases as noted in figure legends. Electron donors (sulfide or thiosulfate) were added to individual bottles from concentrated, anoxic stock solutions for biogenic S(0), sulfide and thiosulfate-only cultures. Other growth conditions, such as light intensity and temperature, remained unchanged.

### Extracellular particle isolation and purification

Extracellular particles were isolated and purified by differential centrifugation (53). One liter of *Cba. tepidum* culture was centrifuged at 10,000 x *g* for 10 min at 4°C in a JA-14 rotor (Beckman Coulter), and the supernatant was centrifuged again at 12,000 x *g* to remove cells and large debris. The supernatant was filtered through a 0.22-μm pore-size polyvinylidene difluoride (PVDF) filter membrane (Millipore, USA). Particles from the filtrate were concentrated by ultracentrifugation at 100,000 x *g* for 2 h at 4°C using an SW32Ti rotor in an Optima LE-80K centrifuge (Beckman Coulter). The pellets from different centrifuge tubes were pooled and resuspended in 1 mL of 1X PBS and pelleted again by ultracentrifugation at 165,700 x *g* for 2 h at 4°C in a TLA-55 rotor (Beckman Coulter). The pellet was resuspended in 500 µL of 1X PBS and stored at −20°C until use. The total protein concentration of purified OMVs was determined using the fluorometric Qubit protein assay (ThermoFisher Scientific) according to the manufacturer’s guidelines.

### Particle characterization by MRPS

Particle samples for quantification from cultures were prepared by centrifugation through a 0.2 μm filter device at 16,000 x *g* for 2 minutes in an Eppendorf 5418 centrifuge. The flow through was used diluted 100-fold with 0.02 µm filtered 1X PBS with 0.1% w/v BSA before analysis. MRPS was performed using an nCS1 instrument (Spectradyne, Torrance, CA, USA) equipped with disposable polydimethylsiloxane cartridges (mold ID 103S and box number 221118). The running buffer was 0.2 µm filtered 1X PBS with 1% (v/v) Tween-20. Diluted OMV samples (7 µL) were loaded, and the particle detection threshold was set after three 10-s acquisitions. Data was collected until the error associated with the concentration measurement was < 3%. The collected data were analyzed with nCS1 Data Analyzer software (Spectradyne). The particle distribution of the dilution buffer was also measured and subtracted from OMV sample data to produce the OMV size and frequency distributions shown. Examples of raw nCS1 particle count data are provided in supplemental materials (Fig. S4).

### Electron microscopic imaging of purified particles

The particle suspension from *Cba. tepidum* was negatively stained with aqueous 2% (w/v) uranyl acetate. Briefly, 400-mesh carbon-coated copper grids (Electron Microscopy Sciences) were glow discharged using a PELCO easiGlow™ glow discharge system (Ted Pella) and then floated on a drop of the vesicle suspension for several seconds. The grids were washed with four drops of water and then negatively stained with aqueous 2% uranyl acetate. Samples were examined in the bioimaging core facility at the University of Delaware using a Zeiss Libra 120 TEM equipped with a Gatan Ultrascan1000 CCD (Fig. 2) or a Thermo Scientific Talos L120C TEM equipped with a Thermo Scientific Ceta-M camera (Fig. 5). For each analysis, at least 15 fields were imaged. To examine particles attached to cells, *Cba. tepidum* culture fixed with 0.4% formaldehyde was centrifuged at 10,000 x *g* for 5 minutes, the supernatant was removed, and the pellet was processed for TEM imaging as described above. Particle size was determined by analysis with ImageJ software (v1.54g). ImageJ was also used to select and scale individual regions of interest from different experiments to the same magnification to compare particle structure across experiments (Fig. S3).

### LPS purification and analysis

LPS was extracted from a suspension of 4×10^10^ particles mL^-1^ by a previously described method (60). LPS was separated by SDS-PAGE using 16% Tricine protein gels (1.0 mm, 14 wells, BioRad) and Tricine SDS running buffer with LPS from *E. coli* O55:B5 as a standard (InvivoGen). The gel was run at 125 V for 120 min at room temperature and stained with Pro-Q Emerald 300 (Thermo Fisher) per the manufacturer’s instructions. Briefly, the gel was oxidized with periodic acid and then washed with 3% glacial acetic to remove residual periodate. The gel was then incubated in 25-fold diluted Pro-Q Emerald 300 staining solution. Gel images were collected on a BioRad ChemiDoc system using Image Lab software version 5.0. Images were automatically contrast adjusted by the software and exported as a TIFF file to produce the figures here.

### Mass spectrometry of lipid A

Mild acid hydrolysis was used to separate lipid A from LPS/LOS by resuspending 0.6 mg of lyophilized purified LPS in 1 mL sodium acetate (10 mM, pH = 4.5) and 1% w/v sodium dodecyl sulfate (SDS). The suspension was boiled (100 ºC, 1h) and then lyophilized. Lyophilized material was washed once with 1 mL of ice-cold acidified ethanol, followed by three washes with ice-cold 95% ethanol to remove SDS from the solid material collected by centrifugation (3,500 x *g*; 4 ºC; 5 min). The final lipid A samples were lyophilized before analysis by mass spectrometry. Samples were reconstituted in 10 uL methanol, then diluted into 5:4:1 chloroform:methanol:water. Lipid A analysis was performed on an UltiMate 3000 UPLC system coupled with an Orbitrap Q-Exactive (Thermo Fisher Scientific) mass spectrometer. A 5 µL sample was injected by syringe at a flow rate of 0.2 mL/min, with the column temperature set at 40 °C. Samples were separated in isocratic mode (50:50 0.1% (v/v) formic acid in LC-MS grade H_2_O:0.1% (v/v) formic acid in LC-MS grade acetonitrile). After separation, the mass spectrometry data were acquired on a Q-Exactive Orbitrap (Thermo Fisher Scientific) equipped with heated electrospray ionization (HESI-II) operating in negative ionization mode. The HESI-II was maintained at a spray voltage of −3.5 kV, and the mass range was set from 600-2200 m/z. Data were processed using Thermo Scientific Xcalibur software version 3.1 (Thermo Fisher Scientific). The lipid A molecular species were identified based on using observed mass to charge (m/z) ratios to search the Lipid MAPS databases (61).

### Quantification of 2-Keto-3-Deoxy-D-Mannooctanoic Acid (KDO)

KDO was derivatized with 1,2-diamino-4,5-methylenedioxybenzene dihydrochloride (DMB, Sigma-Aldrich), and the resulting derivatives were quantified by HPLC following a previously described method (62). Briefly, a KDO standard (Sigma-Aldrich), and purified LPS samples were hydrolyzed using trifluoroacetic acid (TFA). DMB solution (5 mM, 125 mL) prepared according to Hara et al., 1987 (63) was added to hydrolyzed samples, and the mixture was placed in a heat block (50 ºC, 2.5 hrs) covered with aluminum foil. Subsequently, samples were placed on ice (10 min), centrifuged (5 min, 10,000 rpm, 4 ºC), and analyzed using a Shimadzu SCL-40 HPLC system. A Prevail C18 (150 x 4.6 mm, 5 µm particle size, Alltech) analytical column was used; 10 µL of sample was injected; separation was in isocratic mode (86% of 0.1%TFA, 7% Acetonitrile, and 7% Methanol) at 1.3 mL min^-1^ for 20 min. DMB derivatives were detected by fluorescence with excitation at 373 nm and emission at 448 nm.

### Raman Microscopy-Spectroscopy

Samples (40 µL) of S(0) globules or OMVs in 1X PBS were pipetted onto aluminum foil held by an HPLC screw top cap taped to a microscope slide and allowed to dry. Aluminum foil is a non-reactive substrate for Raman measurements. A confocal Raman microscope (LabRam HR Evolution, Horiba Scientific, Kyoto, Japan) was used to record data at the Surface Analysis Facility, University of Delaware. A 532-nm laser was used, and the laser power applied was set at 100%, which corresponds to a laser power of 20 mW on samples. Raman Spectra were obtained over 5 seconds of acquisition time with a wavenumber from 400-4000 cm^-1^ with six accumulations per spectrum.

### Combined Atomic Force Microscopy and Infrared Spectroscopy (AFM-IR)

AFM-IR analysis was performed with a NanoIR2 instrument (Anasys Instruments, Santa Barbara, CA, USA, now Bruker) at the Advanced Materials Characterization Laboratory, University of Delaware. A 5 µL aliquot of samples was spread on a zinc sulfide sampling flat and placed on the stage of the AFM-IR instrument. All AFM-IR measurements and images were done under contact mode. A 1 μm × 1 μm scan area was imaged at a scan rate of 0.6 Hz with a resolution of 384 pixels in both x and y axes. Individual AFM-IR spectra were collected using a MIRcat-2400 laser (Daylight Solutions, San Diego, CA, USA) across the 950-1900 cm^-1^ frequency range. The IR laser produced pulses of 140 ns at the repetition rate of 350 kHz. Analysis Studio software (Anasys Instruments, Santa Barbara, CA, USA) was used for image and spectral data collection.

### S(0) Globule Purification

Cultures (1 L) of strain WT2321 were grown on sulfide-only Pf-7 (4 mM sulfide) in narrow-mouth screw-cap bottles with an open phenolic cap and butyl rubber septa at 30 μmol photon m^-2^ s^-1^ for 36 h. To qualitatively assay for the presence of sulfide, equal volumes of culture supernatant were mixed with 10 mM CuCl_2_, where the formation of a distinct grey precipitate indicated the presence of sulfide greater than 0.2 mM. Sulfide was added to cultures when it was no longer detectable by this assay. Cultures were transferred into sterile 250 ml centrifuge bottles with o-ring sealing caps (Nalgene, Thermo Fisher Scientific). Cells and S(0) were collected by centrifugation (1,000 × *g*, 10 min, 10 °C, JA-14 rotor). The supernatant was removed and cells plus S(0) were suspended in a minimal volume of S-free Pf-7 that was layered over 25 ml of sterile 2.5 M sucrose solution in sterile 50 ml centrifuge tubes with o-ring sealing caps (Nalgene, Thermo Fisher Scientific); at a density of 1.32 g ml^-1^, the sucrose retains cells in the supernatant and selectively pellets S(0) when centrifuged (6,000 × *g*, 50 min, 4 °C, JS-13.1 rotor). The supernatant was removed, and the resuspended pellet was centrifuged through 2.5 M sucrose two more times. The collected S(0) pellet was washed three times to remove sucrose by suspending in S-free Pf-7 and centrifuging (17,500 × *g*, 5 min, 10 °C, JS-13.1 rotor). S(0) was suspended in S-free Pf-7 and immediately distributed into aliquots for characterization studies; aliquots were stored at −80 °C until use.

### Proteomic profiling of OMVs, S(0) globules, and whole cells

Purified OMVs, S(0) globules and whole cell pellets were dissolved in SDS buffer (2% SDS, 50 mM DTT, and 100 mM Tris-HCl pH 8.0) and sonicated for 3 min (pulse: 20s on, 20s off). After sonication, the suspension was incubated at 95 ºC for 10 min. A clear lysate was obtained after centrifugation (16,000 x *g*, 10 min, 25 ºC, Eppendorf 5415 centrifuge). Proteins in the lysate were processed for proteomics analysis following the E3 filter procedure described recently (64). The digests were desalted using C18-based StageTips (CDS Analytical, Oxford, PA). Liquid chromatography-tandem mass spectrometry (LC-MS/MS) was performed on an Ultimate 3000 nanoLC system coupled to an Orbitrap Eclipse Tribrid mass spectrometer with FAIMS Interface (Thermo Scientific). Peptides were separated on a C18 reversed phase column (PepMap100 C18, 25 cm × 75 μm i.d., 3 μm; Thermo Scientific) at a flow rate of 250 nl/min. Mobile phase A consisted of 0.1% formic acid in water. Mobile phase B consisted of 0.1% formic acid in acetonitrile. A gradient separation was used: mobile phase B was held at 1% for 5 min, increased to 25% over 90 min, reached to 35% at 110 min, increased to 80% over 10 min, then re-equilibrated at 1% mobile phase B for 15 min. For data-dependent MS acquisition, the spray voltage was set to 1.8 kV, funnel RF level at 50%, and heated capillary temperature at 275 °C. The MS data were acquired in Orbitrap mode at 60,000 resolution, followed by MS/MS acquisition of the most intense precursors for 1 s. For MS2 analysis, collision was set by HCD at 30% normalized collision energy (NCE). For FAIMS settings, a 3-CV experiment (−40|-55|-75) was applied.

Peptide identification and quantitation used MaxQuant and Andromeda software (version 1.6.3.4) with most default settings. *Cba. tepidum* proteome database (Taxon ID 194439; 2,251 sequences) was obtained from UniProt Knowledgebase. The enzyme specificity was set to ‘Trypsin’; oxidation of methionine, and acetyl (protein N-terminus) were set as variable modification; carbamidomethylation of cysteine was set as fixed modification. Up to 2 missed cleavages were allowed. The false discovery rate (FDR) was set to 1% for protein and peptide identifications. MaxLFQ function embedded in MaxQuant was enabled for label-free quantitation, and the LFQ minimum ratio count set to 1.

### Bioinformatic classification of identified proteins

Visualization of quantitative proteomic data was mainly performed using Perseus software (version 1.6.2.3) (65). Label-free quantitation intensities generated by MaxQuant were log2-transformed and imported into Perseus with protein annotations. As noted in the Results, the thiosulfate OMV samples were removed, and data were filtered to only contain proteins observed in at least 2/12 samples (n = 1,447 unique proteins). Uniprot accessions for proteins associated with each sample type were exported for to FunRich v. 3.1.3 (66) to construct a Venn diagram (Fig. 4A). NaNs were replaced using randomly selected values from a normal distribution with the default settings (downshift value = 1.8 standard deviations, width = 0.3). The NaN replaced data were used for principal component analysis (PCA). The PCA scores and loadings for each protein were exported from Perseus to Excel to produce the plots (Fig. 4B and 4C) The subcellular localization of each protein was predicted using the PSORTb subcellular localization tool version 3.0.3 (https://www.psort.org/psortb/) and merged with the data from Perseus in Excel to color loading values of individual proteins based on predicted localization (Fig. 4B). The Perseus session file and Excel file containing all data in Figure 4 are available at: https://hansonlabgit.dbi.udel.edu/hanson/ctep_omv_manuscript

### S(0) Quantification in OMVs

Three independent 20 mL cultures of *Cba. tepidum* were grown with 7 mM sulfide for 14 hrs. OMVs were purified from the entire volume of each culture as above and resuspended in 20 microL of 1X PBS after the second ultracentrifugation. A 15 microL aliquot of OMV suspension was extracted with 1 mL of hexanes (Sigma Aldrich). S(0) was quantified in the hexane phase by HPLC on a 4.6 x 150 mM Prevail C18 column eluted with 90% methanol at 1 mL min^-1^ following absorbance at 280 nm on a Shimadzu SCL-40 HPLC system. Sulfur powder (Sigma-Aldrich) dissolved in chloroform was diluted to known concentrations in hexane to generate a standard curve. The remaining OMV suspension was used for TEM imaging as described above.

## ACKNOWLEDGMENTS

This research was partially supported by a National Science Foundation research grant to TEH (1244373). VLN was partially supported by a grant from the W.M. Keck Foundation (16A01455). ATL was supported by an NSF Graduate Research Fellowship Program award (0750966). Research reported in this publication was supported by the National Institute of General Medical Sciences of the National Institutes of Health under Award Numbers T32GM133395 (partial support for VLN) and P20GM104316 for the Orbitrap Eclipse MS instrument. Microscopy equipment was acquired with Unidel Foundation support and a Major Research Instrumentation award (1828325) for the purchase of the LabRam HR Evolution. Microscopy access was supported by grants from the NIH-NIGMS (P20GM103446), the NIGMS (P20GM139760) and the State of Delaware. The content is solely the responsibility of the authors and does not necessarily represent the official views of the W.M. Keck Foundation, the National Institutes of Health, or the National Science Foundation.

