## Supplemental Methods-Figures-Tables for "*Chlorobaculum tepidum* Outer Membrane Vesicles May Transport Biogenic Elemental Sulfur"

### SUPPLEMENTAL MATERIAL

#### METHODS AND MATERIALS

##### Protein Extraction and Shotgun Proteomic Analysis of *Cba. tepidum* S(0) Globules

A comprehensive analysis of extraction conditions (1) developed the protein extraction method used here. S(0) globules were suspended in 100 mM triethylammonium bicarbonate (TEAB) with 2% w/v CHAPS detergent followed by sonication on ice (2 min increments, 10 mins total) and centrifugation (16,000 × g, 30 mins) to pellet S(0). The supernatant was removed and treated with DetergentOUT$^{TM}$ spin columns (G-Biosciences) followed by concentration in a SpeedVac$^{TM}$ at 37 °C (Thermo Fisher Scientific Inc.). The dried pellet was resuspended in 0.5 M TEAB with 0.1% w/v SDS and reduced by treatment with tris-(2-carboxyethyl)phosphine (TCEP, 4.3 mM, 60 °C, 1 hr). Free cysteines were blocked with methyl methane-thiosulfonate (8.3 mM, 10 mins, room temperature). Proteins were digested overnight (37 °C) with 1 µg trypsin (Promega) per 10 µg protein. Digestion was stopped by adding formic acid to 1.5% v/v. Peptides were cleaned up prior to LC-MS/MS using Bond Elut OMIX C18 tips (Agilent) according to manufacturer instructions, and concentrated to dryness using a SpeedVac$^{TM}$ vacuum concentrator at 37 °C. Dried peptides were stored at -80 °C.

Peptides were separated by low pH reverse phase high performance liquid chromatography on a Tempo LC-MALDI spotter (Eksigent). Peptides were resolubilized in Mobile Phase A (2% acetonitrile, 0.1% trifluoroacetic acid (TFA), loaded onto a 1.2 µl CapRod 18E capillary column (Merck KGaA), and washed with 10 column volumes of 98% Mobile Phase A/2% Mobile Phase B (98% acetonitrile, 0.1% trifluoroacetic acid). Peptides were eluted by an 80 min gradient to 55% acetonitrile and 0.1% TFA. Eluate was spotted in 10 second increments onto stainless steel target plates with α-cyano-4-hydroxycinnamic acid (Sigma-Aldrich) as matrix.

Peptides were identified by MALDI-TOF/TOF mass spectrometry in positive ion reflector mode on an AB Sciex 5800 MALDI-TOF/TOF Analyzer. MS data were collected over a mass range of 800-4000 m/z and were processed with internal calibration. A maximum of 15 precursors, with signal-to-noise ratios of at least 20, were selected per spot for MS/MS. Fragmentation was induced with 1 kV collision energy. MS/MS spectra were processed with default calibration and submitted for Mascot v2.4 database searchers through Protein Pilot software v4.5 (ABSciex). Spectra were searched against an NCBI database of the *C. tepidum* proteome (downloaded October 29, 2014). Peptide identifications with 95% confidence or greater and protein identifications containing at least one significant unique peptide were accepted.

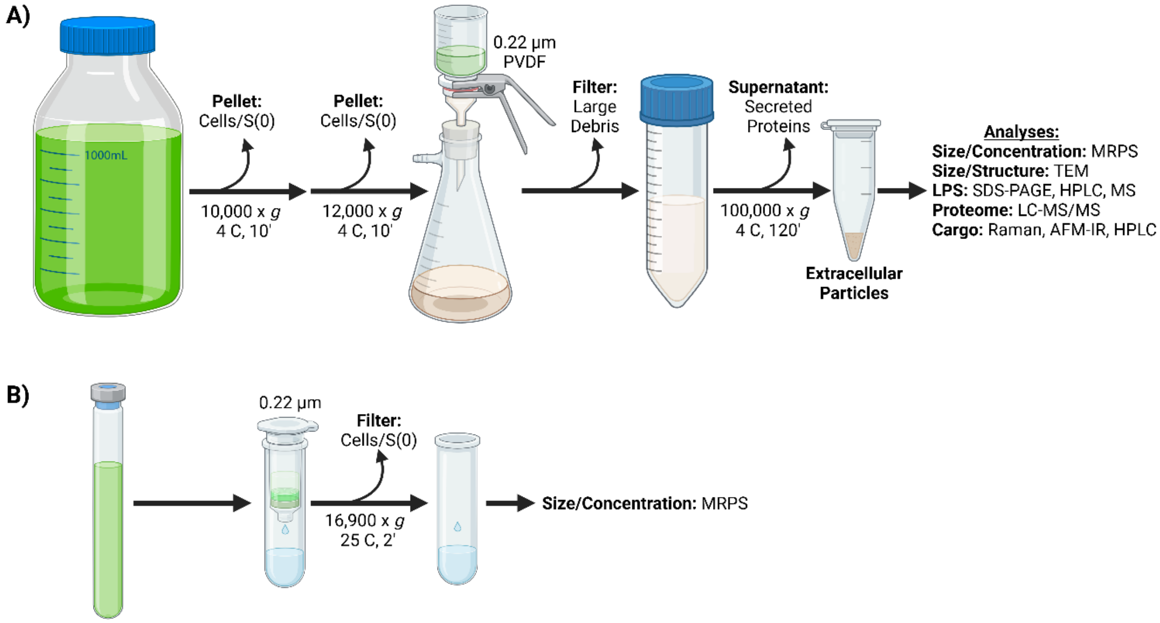


**Figure S1**. Workflow for particle purification methods from *Cba. tepidum* cultures. A) Extracellular particles for analyses requiring large amounts of input material were prepared by differential centrifugation. B) Extracellular particle samples for routine counting and sizing were prepared by centrifugation through a 0.22 µm filter. Non-standard abbreviations: MRPS-microfluidic resistive pulse sensing; AFM-IR-atomic force microscopy with Infrared spectroscopy. Created in BioRender https://BioRender.com/q133wb4

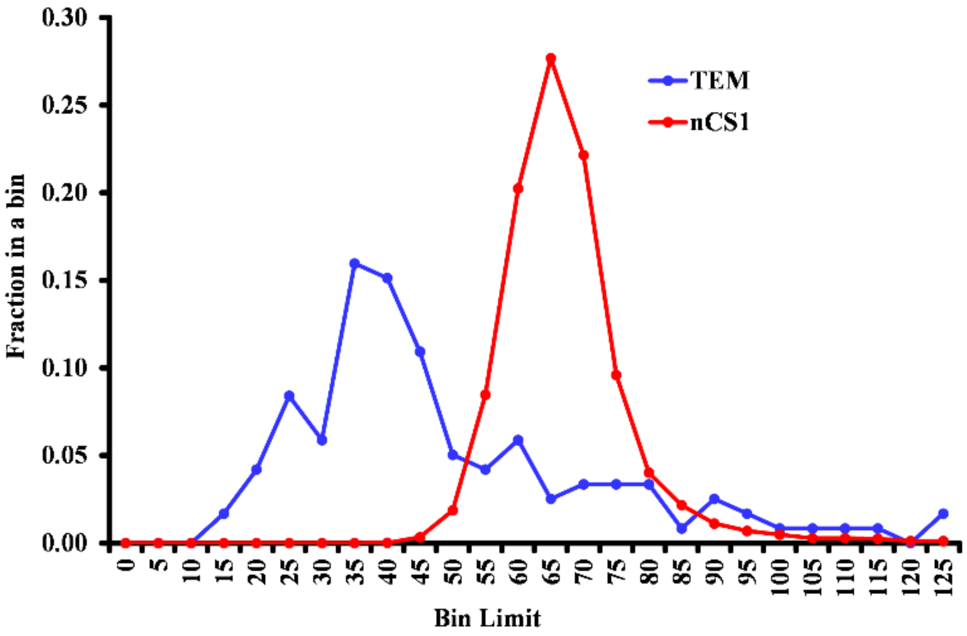


**Figure S2**. Particle size distributions by TEM and MRPS

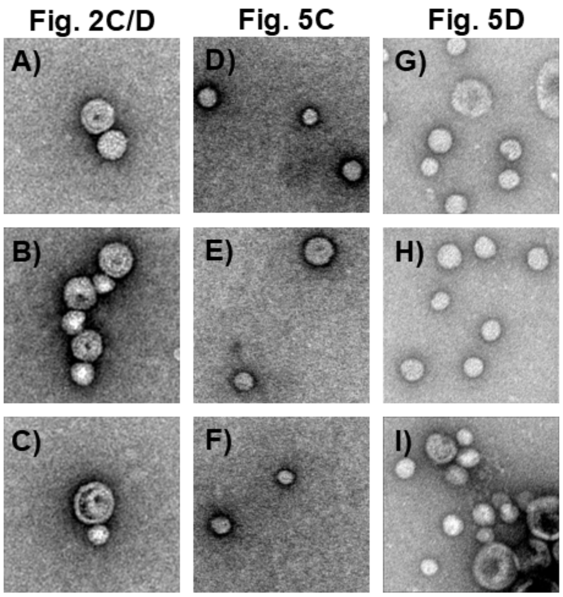


**Figure S3**. Close up images of OMVs from TEM images used in Figures 2 (A-C) and 5 (D-I). Each image in this collection is 200 nm on each side and were produced by selecting, copying and scaling regions of interest from the initial images in ImageJ. No changes were made to any other image parameters.

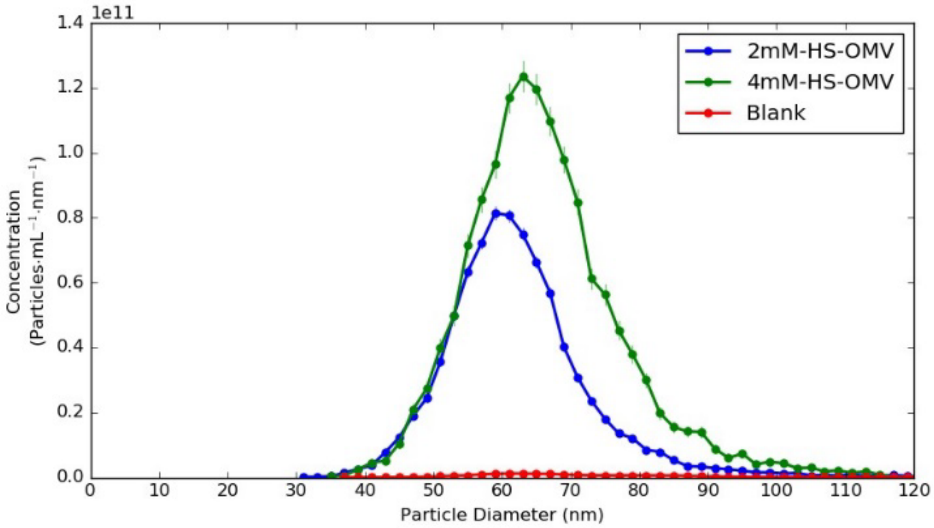


**Figure S4**. Example of raw nCS1 particle count data

**Table S1**. Proteins identified by shotgun proteomic analysis of S(0) globules produced by wild type *Cba. tepidum*. Proteins also identified in our gel-based studies (1, 2) or in OMV proteomic studies from the literature are indicated.

| Protein Information | | | | | | Detected in: | | | |
|---|---|---|---|---|---|---|---|---|---|
| Locus Tag | Accession[1] | Gene Name | Description | Predicted Location [2] | COG Cat.[3] | S(0) shotgun | S(0) gel | Cells gel | OMVs Reference |
| CT0018 | NP_660924 | *atpH* | ATP synthase F0F1 subunit delta | C | C | X | | | 10 |
| CT0020 | NP_660926 | *atpE* | ATP synthase F0 subunit C | CM# | C | X | | | 9 |
| CT0027 | NP_660933 | | hypothetical protein CT0027 | U | -- | X | | | |
| CT0031 | NP_660937 | *ftsA* | cell division protein FtsA | C | D | X | | | 9,10 |
| CT0068 | NP_660974 | | hemagglutinin-related protein | OM# | M | | X | | 5,7 |
| CT0089 | NP_660995 | *clpB-2* | ATP-dependent Clp protease, ATP-binding subunit ClpB | C | O | X | | X | 3,9,10 |
| CT0131 | NP_661037 | | hypothetical protein CT0131 | C | R | X | | | |
| CT0150 | NP_661056 | *nusG* | transcription antitermination protein NusG | C | K | X | | | 3,9 |
| CT0153 | NP_661059 | *rplJ* | 50S ribosomal protein L10 | C# | J | X | | | 4,9,10,12 |
| CT0154 | NP_661060 | *rplL* | 50S ribosomal protein L7/L12 | U | J | X | | | 4 |
| CT0155 | NP_661061 | *rpoB* | DNA-directed RNA polymerase subunit beta | C | K | X | | X | 3,4,9-11 |
| CT0156 | NP_661062 | *rpoC* | DNA-directed RNA polymerase subunit beta' | C | K | X | | | 3,4,9-11 |
| CT0159 | NP_661065 | *efp* | elongation factor P | C | J | X | | | 9 |
| CT0160 | NP_661066 | *hupB* | DNA-binding protein HU-beta | C | L | X | X | X | 9-12 |
| CT0173 | NP_661079 | *serB* | phosphoserine phosphatase | C | E | X | | | 11 |
| CT0249 | NP_661153 | | glutathione S-transferase | C | E | X | | | |
| CT0254 | NP_661158 | *ompH* | outer membrane protein OmpH | U | M | X | | | 4,5,13 |
| CT0264 | NP_661168 | *rho* | transcription termination factor Rho | C | K | X | | | 10 |
| CT0288 | NP_661192 | *rpsA* | 30S ribosomal protein S1 | C | J | X | | | 3,9,10 |
| CT0302 | NP_661206 | *petC5* | cytochrome b6-f complex, iron-sulfur subunit | CM | C | X | X | X | 3 |
| CT0303 | NP_661207 | *petB* | cytochrome b-c complex, cytochrome b subunit | CM | C | X | | | 3,9,10 |
| CT0312 | NP_661216 | | DnaK suppressor protein | C | T | X | | | |
| CT0350 | NP_661254 | *fabI* | enoyl-(acyl-carrier-protein) reductase | CM | I | X | | | 3,12 |
| CT0529 | NP_661429 | *groS* *groES* | co-chaperonin GroES | C | O | X | | X | 9 |
| CT0530 | NP_661430 | *groL* *groEL* | chaperonin GroEL | C | O | X | X | X | 3-5,8-13 |
| CT0531 | NP_661431 | | sensor histidine kinase/response regulator | CM | T | X | | | 10,11 |
| CT0547 | NP_661447 | *mreB-1* | rod shape-determining protein MreB | C | D | X | | | 3,5,10 |

| | | | | | | | | | |
|---|---|---|---|---|---|---|---|---|---|
| CT0563 | NP_661463 | *tyrS* | tyrosyl-tRNA synthetase | C | J | X | | | |
| CT0607 | NP_661507 | | hypothetical protein CT0607 | C | S | X | | | |
| CT0638 | NP_661535 | | peptidoglycan-associated lipoprotein | OM | M | X | | | 3,4,9-11,13 |
| CT0642 | NP_661539 | | hypothetical protein CT0642 | C | -- | X | | X | |
| CT0643 | NP_661540 | *dnaK* | molecular chaperone DnaK | C | O | X | X | X | 5,8-10 |
| CT0644 | NP_661541 | | HSP20 family protein | C | O | X | X | X | |
| CT0829 | NP_661723 | *htpG* | heat shock protein 90 | C | O | X | | X | 9,10 |
| CT0841 | NP_661735 | *trx-2* | thioredoxin | C | CO | X | | X | |
| CT0893 | NP_661786 | | hypothetical protein (Porin_5) | OM | -- | X | X | | 11 |
| CT0903 | NP_661796 | | transcriptional regulator | C | P | X | | | |
| CT0941 | NP_661834 | *btuR, CobO, CobP* | cob(I)alamin adenosyltransferase | C | H | X | | | |
| CT0960 | NP_661853 | *purC* | phosphoribosylaminoimidazole-succinocarboxamide synthase | C | F | X | | | 9,10 |
| CT0980 | NP_661873 | | ArsA ATPase | C | P | X | | | |
| CT1007 | NP_661900 | | hypothetical protein CT1007 (DsrE/DsrF - like family) | CM | -- | X | | | |
| CT1054 | NP_661945 | *prc* | carboxyl-terminal protease | CM | M | X | | | 4,7-11 |
| CT1133 | NP_662024 | | hypothetical protein (CRISPR-associated protein) | U | -- | | X | | |
| CT1170 | NP_662061 | | hypothetical protein CT1170 | C | I | X | | | 9 |
| CT1225 | NP_662115 | | N-acetylmuramoyl-L-alanine amidase | U | V | X | | | 9-11 |
| CT1239 | NP_662127 | *secA* | preprotein translocase subunit SecA | C | U | X | | X | 3,9-11 |
| CT1297 | NP_662185 | *bchI chlI* | magnesium-chelatase subunit I | C | R | X | | | |
| CT1309 | NP_662197 | | hypothetical protein CT1309 | U | -- | X | | | |
| CT1353 | NP_662240 | | OmpA family protein | OM | M | X | | X | 4,9-13 |
| CT1361 | NP_662248 | *prs prsA* | ribose-phosphate pyrophosphokinase | C | EF | X | | | 3,4,9,10 |
| CT1362 | NP_662249 | *rplY ctc* | 50S ribosomal protein L25 general stress protein | C | J | X | | | 9,10 |
| CT1447 | NP_662333 | | serine protease | P | O | X | X | X | 3,4,10,11,13 |
| CT1485 | NP_662370 | *grpE* | heat shock protein GrpE | C | O | X | | | 9 |
| CT1499 | NP_662384 | *fmoA* | bacteriochlorophyll A protein | Csm | -- | X | | X | |
| CT1577 | NP_662460 | *frr* | ribosome recycling factor | C | J | X | | X | 9 |
| CT1591 | NP_662474 | *ribBA* | 3,4-dihydroxy-2-butanone 4-phosphate synthase | C | H | X | | | 3 |
| CT1649 | NP_662532 | *pnp* | polynucleotide phosphorylase/polyadenylase | C | J | X | | | 3,5,9,10,12 |
| CT1742 | NP_662622 | *feoB-1* | ferrous iron transport protein B | CM | P | X | | | |

| | | | | | | | | | |
|---|---|---|---|---|---|---|---|---|---|
| CT1743 | NP_662623 | *feoA-1* | ferrous iron transport protein A | U | P | X | | | |
| CT1744 | NP_662624 | | hypothetical protein CT1744 | U | P | X | | | |
| CT1745 | NP_662625 | | hypothetical protein CT1745 | U | -- | X | | X | |
| CT1780 | NP_662659 | *tsf* | elongation factor Ts | C | J | X | | | 3,9,12 |
| CT1781 | NP_662660 | *rpsB* | 30S ribosomal protein S2 | C | J | X | | | 3,4,10,11,13 |
| CT1782 | NP_662661 | *rpsI* | 30S ribosomal protein S9 | C | J | X | | | 3,4,10,11,13 |
| CT1785 | NP_662664 | | ATP-binding Mrp/Nbp35 family protein | CM | D | X | | | 3,4,10 |
| CT1804 | NP_662683 | | hypothetical protein | OM | -- | X | X | X | |
| CT1833 | NP_662712 | *gatC* | aspartyl/glutamyl-tRNA amidotransferase subunit C | C | J | X | | | 3 |
| CT1867 | NP_662744 | | hypothetical protein CT1867 | P | S | X | | | |
| CT1921 | NP_662798 | | cysteine synthase/cystathionine beta-synthase | C | E | X | | | 8,9 |
| CT1939 | NP_662816 | | ArsA ATPase | C | P | X | | | |
| CT1942 | NP_662819 | *csmA* | chlorosome envelope protein A | Csm | -- | X | | X | |
| CT1943 | NP_662820 | *csmC* | chlorosome envelope protein C | Csm | -- | X | X | X | |
| CT1947 | NP_662824 | *ssb-1* | single-strand binding protein | C | L | X | | | 6,9 |
| CT1955 | NP_662832 | | magnesium-chelatase, bacteriochlorophyll c-specific subunit | C | H | X | | X | |
| CT1970 | NP_662846 | | HSP20 family protein | U | O | X | X | X | 10 |
| CT1986 | NP_662862 | | hypothetical protein CT1986 | U | R | X | | | |
| CT2001 | NP_662877 | *bcp-2, ahpC, tsa* | bacterioferritin comigratory protein, thiol peroxidase | C | O | X | | | 8,9 |
| CT2026 | NP_662901 | | c-type cytochrome | U | -- | X | | | |
| CT2033 | NP_662908 | *atpA* | ATP synthase F0F1 subunit alpha | C | C | X | | X | 3,5,8-10 |
| CT2047 | NP_662922 | | AcrB/AcrD/AcrF family protein | CM | V | X | | | 10 |
| CT2049 | NP_662924 | | LipD protein, putative | OM | MU | X | X | | 4,11,13 |
| CT2054 | NP_662929 | *csmB* | chlorosome envelope protein B | Csm | -- | X | | X | |
| CT2067 | NP_662942 | | pentapeptide repeat-containing protein | E | S | X | | | |
| CT2097 | NP_662971 | | hypothetical protein CT2097 | U | S | X | | | |
| CT2101 | NP_662975 | | hypothetical protein CT2101 | C | S | X | | X | |
| CT2129 | NP_663003 | *rplT* | 50S ribosomal protein L20 | C | J | X | | | 4,9,10,12 |
| CT2144 | NP_663018 | | outer surface protein, putative | OM | M | X | | X | 5,12 |
| CT2147 | NP_663021 | | hypothetical protein CT2147 | C | -- | X | | | |
| CT2151 | NP_663025 | *bchB* | light-independent protochlorophyllide reductase subunit B | C | C | X | | | |
| CT2160 | NP_663034 | *gidB rsmG* | 16S rRNA methyltransferase GidB | C | J | X | | | |

| | | | | | | | | |
|---|---|---|---|---|---|---|---|---|
| CT2161 | NP_663035 | *rplQ* | 50S ribosomal protein L17 | C | J | X | X | 4,9,10,12 |
| CT2162 | NP_663036 | *rpoA* | DNA-directed RNA polymerase subunit alpha | C | K | X | | 3,9,10,12 |
| CT2177 | NP_663051 | *rplE* | 50S ribosomal protein L5 | C | J | X | | 3,4,9,10,12,13 |
| CT2182 | NP_663056 | *rplP* | 50S ribosomal protein L16 | C | J | X | | 4,9,12 |
| CT2186 | NP_663060 | *rplB* | 50S ribosomal protein L2 | C | J | X | | 3,4,9,10,12,13 |
| CT2191 | NP_663065 | *tuf* | elongation factor Tu | C | J | X | | 3,4,7-11,13 |
| CT2215 | NP_663089 | *gatB* | aspartyl/glutamyl-tRNA amidotransferase subunit B | C | J | X | | 3,8 |
| CT2216 | NP_663090 | | hypothetical protein | CM | S | X | | |
| CT2234 | NP_663108 | *atpD-2* | ATP synthase F0F1 subunit beta | C | C | X | X | 3,5,8-10 |
| CT2264 | NP_663137 | *surA, prsA* | peptidyl-prolyl cis-trans isomerase SurA | P | O | X | | 4,9,10,13 |
| CT2281 | NP_663152 | *clpB-1* | ATP-dependent Clp protease, ATP-binding subunit ClpB | C | O | X | | 3,9,10 |

[1] RefSeq (NP_xxxxxx) or GenBank (AAYxxxxx) Accession

[2] Location prediction by PSORTb, except for except for manual annotations noted with #. Csm: chlorosome; C: cytoplasmic; CM: cytoplasmic membrane; OM: outer membrane; P: periplasmic; U: unknown.

[3] Abbreviations for COG functional categories are provided in Table S2.

**Table S2.** Functional classification of proteins associated with S0 based on COG categories, and portion of proteins within COG categories with homologs previously identified in proteomic studies of outer membrane vesicles.

| COG Category | | $S^0$ proteins | | $S^0$ proteins with OMV protein homologs | |
|---|---|---|---|---|---|
| | | count[1] | % of total[1] | count | % in COG category |
| C | Energy production and conversion | 8 | 7.9% | 6 | 75% |
| D | Cell cycle control, cell division, chromosome partitioning | 3 | 3.0% | 3 | 100% |
| E | Amino acid metabolism and transport | 4 | 4.0% | 3 | 75% |
| F | Nucleotide metabolism and transport | 2 | 2.0% | 2 | 100% |
| H | Coenzyme metabolism and transport | 3 | 3.0% | 1 | 33% |
| I | Lipid metabolism and transport | 2 | 2.0% | 2 | 100% |
| J | Translation, ribosomal structure and biogenesis | 20 | 19.8% | 18 | 90% |
| K | Transcription | 5 | 5.0% | 5 | 100% |
| L | Replication, recombination and repair | 2 | 2.0% | 2 | 100% |
| M | Cell wall/membrane/envelope biogenesis | 7 | 6.9% | 7 | 100% |
| O | Post-translational modification, protein turnover, chaperones | 13 | 12.9% | 11 | 85% |
| P | Inorganic ion transport and metabolism | 6 | 5.9% | 0 | 0% |
| R | General Functional Prediction only | 3 | 3.0% | 0 | 0% |
| S | Function Unknown | 6 | 5.9% | 0 | 0% |
| T | Signal Transduction mechanisms | 2 | 2.0% | 1 | 50% |
| U | Intracellular trafficking, secretion, and vesicular transport | 2 | 2.0% | 2 | 100% |
| V | Defense Mechanisms | 2 | 2.0% | 2 | 100% |
| | No COG assigned | 14 | 13.9% | 1 | 7% |

[1] Count and % sum to greater than 101 and 100%, respectively, as a small number of proteins were assigned to more than one COG category.
